# Serological evidence of Rift Valley fever infection and risk factors among one-humped camels (*Camelus dromedarius*) in Northern Nigeria

**DOI:** 10.1101/2020.01.10.901470

**Authors:** Adamu Andrew Musa, Yila Simon Ayo, Allam Lushakyaa, Sackey Anthony, Alhaji Nma Bida, Garba Bello Sikiti, Mambula-Machunga Salamatu, Nafarnda Wesley Daniel, Idoko Sunday Idoko, Balogun Oluwadare Emmanuel, Owolodun Olajide Adewale, Dzikwi Asabe Adamu

## Abstract

**Background:** Rift Valley fever (RVF) is a zoonotic disease that has become emerging and re-emerging in some regions of the world, infecting livestock and humans. One-humped camels are important economic livestock species in Africa used for traction, transportation, and food. Regional and international trade has continued to increase the risk of this disease, spreading widely and causing severe economic and public health catastrophes in affected regions. In spite of these risks, there is a dearth of information about the status of RVF in camels in Nigeria. This study was carried out to determine the prevalence of the RVF virus in one-humped camels in Nigeria and identify the risk factors associated with the disease.

**Methods:** A cross-sectional study with simple random sampling was carried out in seven local government areas of Jigawa and Katsina States. The sera from camels were tested for anti-RVFV IgG. Camel owners were administered a structured questionnaire to ascertain their knowledge, attitude, and practice.

**Results:** An overall prevalence of 19.9% (95% CI; 17.07-22.90) was recorded. Based on age groups, the highest prevalence of 20.9% (95% CI; 17.00-25.31) was obtained among older camels (6-10 years), while female camels recorded a high prevalence of 20.4% (95%CI; 15.71-25.80). Sule Tankar-kar recorded the highest prevalence with 33% (95%CI; 1.31-4.72, p= 0.007) and OR 2.47 in Jigawa State while Mai’adua had 24.7% (95%CI; 0.97-2.73, p=0.030) with OR 1.62 in Katsina State respectively. From the risk map, local government areas bordering Niger Republic were at a high risk of RVF. Only high rainfall was not significantly linked with RVF occurrence among nomadic camel pastoralists (95%CI 0.93-5.20; p=0.070).

**Conclusion:** There is a need for the country to have quarantine units across borders for screening animals coming from neighbouring countries for transboundary infectious diseases such as RVF.

**Author Summary:** Rift Valley fever is a viral haemorrhagic fever that affects animals and humans with high mortality. Recently there has been increased demand in camel meat and products for food and therapeutic purposes. Climate change, coupled with insecurity in the Sahel, has had a significant impact on transhumance activities where camels and their owners move to different countries in search of pasture for their animals. Though Nigeria has not reported an outbreak of Rift Valley fever despite serological evidence in various animal species, there is a need to assess RVF in camels, which is a critical animal species, involved in transhumance with the potential of introducing transboundary diseases into new areas. The study assessed the presence of antibodies in camels, identified risk factors associated with the disease in camels and areas at risk for the disease. Our study found a seroprevalence of 19.9% in camels in two northern states of Nigeria, which shares a boundary with the Niger Republic that recently reported an outbreak. Our findings suggest that areas in proximity to Niger Republic are at a high risk to the disease and camels belonging to transhumance pastoralists are highly likely to contract Rift Valley fever since they are exposed to various ecological and environmental factors that precipitate the disease.

## Introduction

RVF is an arboviral hemorrhagic disease of domestic, wild animals and man[1], caused by the RVF Virus (RVFV), a member of the Genus *Phlebovirus* and Family *Phenuiviridae* [2], a tripartite, negative sense single stranded RNA genome comprising of large (L), medium (M) and small (S) segments [3]. Epidemics have been reported in several African countries with negative impact on the socio-economic livelihood of the affected communities [4,5]. RVF is considered an endemic disease in sub Saharan Africa [6]. In September 2000, an outbreak was reported off the shores of Africa in the Arabian Peninsula and Yemen [7]. Recent outbreaks were recorded in Kenya, Sudan, Uganda and South Africa involving ruminants and humans [8]. Chevalier and collaborators [9] had feared that the disease could find its way to Europe and other parts of the world where an outbreak has never been reported thereby causing a devastating effect. In Nigeria, the disease was first described among Merino sheep imported from South Africa to Vom, Plateau State [10]. The virus was isolated from field population of *Culicoides* and *Culex antennatus* at a farm in Nigeria [11]. RVF virus infection in animals and humans occurs mainly through bite of infected mosquitoes (*Aedes* and *Culex*) and other hematophaguos arthropods or through direct contact with infected animal tissues or fluids by humans due to occupational hazards [12].

Movements of viremic animals along trade and cattle routes have been suspected to be responsible for the spread of RVF [13]. Infection is usually associated with high neonatal mortality and abortions in ruminants with fever, nasal discharge, listlessness, fetid diarrhea could be observed. Infections in humans are characterized by febrile illness, followed by hemorrhagic fever, encephalitis [14,15].Surveillance for RVF in most African countries including Nigeria is limited and outbreaks may go unnoticed and therefore misdiagnosed and under reported. The aim of this study was to search for evidence for circulation of RVFV among camels in Northern Nigerian States that share boundary with Niger Republic.

## Materials and Methods

### Ethics Statement

Research Statement was approved by the University of Abuja Ethical Committee Animal Care Unit (ref UAECAU/2018/025). Camel owners were obtained on the objectives of the study and verbal consent was obtained before the commencement of the study.

### Study design

A cross- sectional study with simple random sampling was carried out from November 2016 to April 2017. Seven hundred and twenty camels from semi arid and arid local government areas of Jigawa and Katsina States were sampled. The camels were properly restrained in a crouching position and 5 ml of blood was collected using a 10 ml syringe with an 18G needle via the jugular vein. Age, sex and location of herd were recorded. Blood was gently transferred into a non anti-coagulant sample bottles and labeled appropriately. Samples were centrifuged at 10,000 g for fifteen minutes to allow proper separation of serum from the clotted blood. Sera were harvested using a sterile pipette into 2 ml cryovial tubes, labeled and stored at − 20°C for further use.

**Fig 1:**
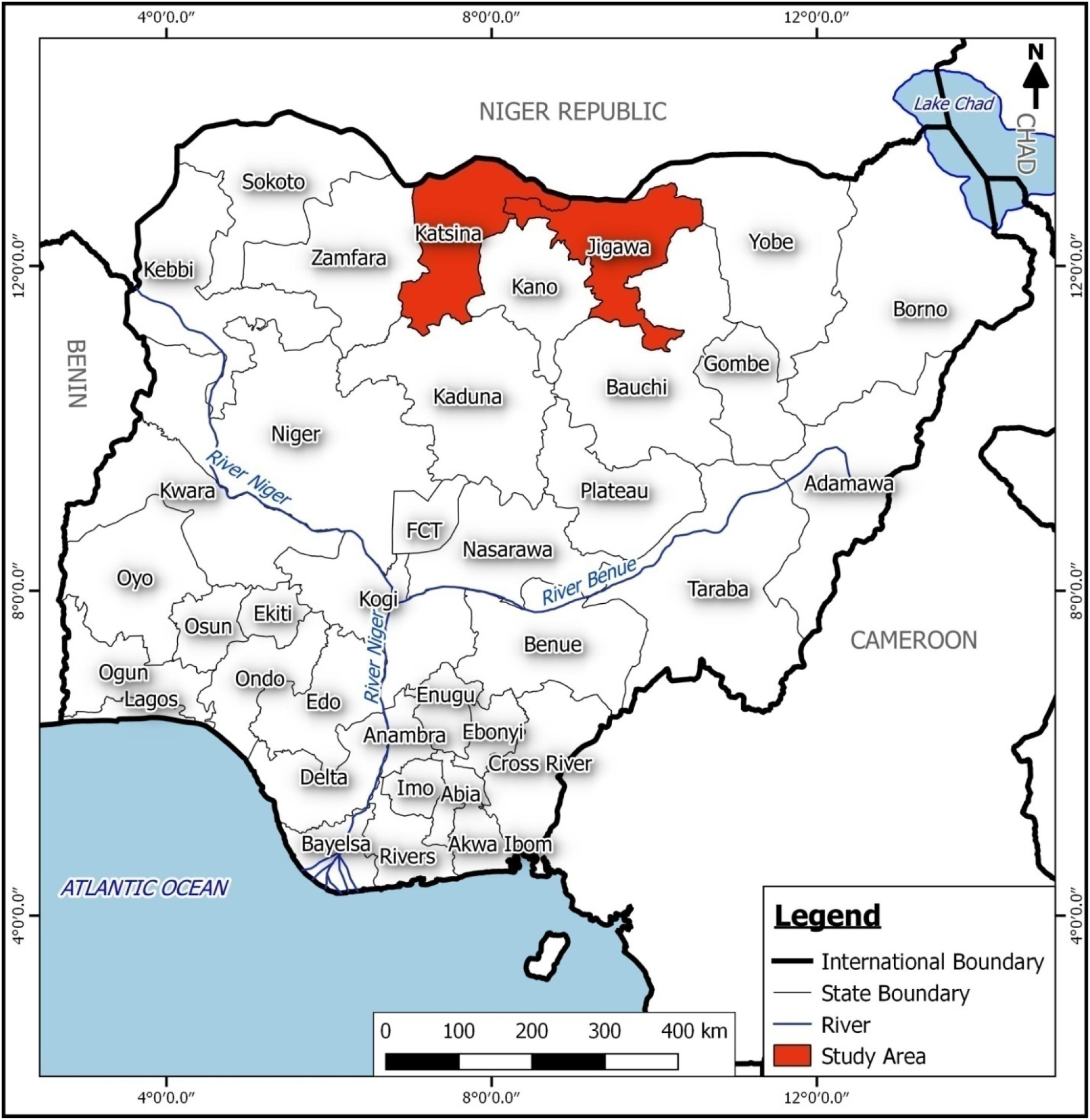
Map of Nigeria showing the study States

### Laboratory Analysis

#### Detection of RVF Virus antibodies using captured IgG ELISA

To determine the past exposure of RVF virus infection, serum samples were tested for anti-RVF virus IgG antibody using multi species competitive ID Screen® RVF IgG ELISA kits (IDVet Innovative Diagnostics, Grabels, France) as described by [16]. The test was validated when the mean value of the positive control O.D. (OD_pc_) was less than 0.3 of the OD_NC_ and the mean value of the negative control O.D. (OD_NC_) is greater than 0.7. The results were read using an ELISA reader set at 450nm. For each sample, a competition percentage (SN %) was calculated using S/N%=OD_sample_/OD_NC_*100. Samples with competition percentage (S/N %) less than or equal to 40% were considered ‘positive’ while those between 40% and 50% were considered ‘doubtful’ and samples with competition percentage above 50% were considered ‘negative’.

### Geospatial Distribution of RVF

Mapping and determination of the geospatial distribution of the occurrence of RVFV in Jigawa and Katsina State was carried out using the ArcGIS software (Environmental Systems Research Institute, Inc; Redlands, USA) [17]. Data on the occurrence of RVFV and the coordinates of the selected Local GovernmentAreas: Maigatari (E9.4381°, N12.81723°); Suletankarka (E9.22928°, N12.66941°); Gumel (E9.38379°, N12.62759°); Babura (E9.01053, N12.76655°) in Jigawa State, while Maiadua (E8.22888°, N13.17292°); Daura (E8.316148°, N13.032475°) and Jibiya (E7.26506°, N13.12238°) (Fig 4.1) in Katsina State were captured with e Trex Legend H version 3.20 global positioning system (GPS) and were imported into Geographical Information System (GIS) software ArcGIS 9.3 (ESRI, USA) to produce disease risk map of the study areas (Fig 4.2).

### Questionnaire design and data collection

Eighty-eight structured questionnaires were administered to camel owners in Katsina States and in Jigawa States respectively to assess their knowledge, attitude, and practice regarding Rift Valley fever. Survey questionnaires asked participants on their socio-demographics, which included age, gender, occupation, and level of education coupled with the knowledge of some clinical signs associated with RVF that includes anorexia, listlessness in newborn, abortion in pregnant animals, hemorrhages, fetid diarrhea, zoonosis, and endemicity. Also, participants were asked on practices employed to mitigate the scourge of RVF, ranging from the use of repellent against arthropods, avoiding swampy areas during grazing, avoiding contact with aborted fetuses, separation of healthy animals from infected animals, loaning of animals and the use of ethnoveterinary practices. Finally, some environmental risk factors were assessed such as high mosquito density, rainfall, irrigated rice fields or dams, bushy vegetation and access to the presence of water bodies

### Statistical Analysis

Results obtained from serological tests were subjected to analysis by Statistical Product and Service Solution (SPSS) version 23. Descriptive statistics was used to test for association between categorical variables with p-value ≤ 0.05 considered significant. Participants’ responses were first summarized into Microsoft Excel 7 (Microsoft Corporation, Redmond, WA, USA) spreadsheets. Data were analyzed using OpenEpi version 2.1. Frequencies and proportions were used for descriptive analysis. Categorical variable responses were presented as proportions and their associations determined by logistic regression models. To assess the association of factors that influence RVF emergence in camel settlements, the factors were considered as independent (explanatory) variables, while respondents' overall response levels constituted the dependent (outcome) variables. However, to create outcome variables, a unique scoring system was designed for the responses. Each respondent was assigned a response score within a range of 1– 20 points and converted to 100%. These scores reflected the stringency of their responses to questions. The score range was further categorized into ‘poor’ or ‘satisfactory’ to keep them as binary variables. Response scores that fell within 1–10 points were considered ‘poor’ (≤49%), and those that fell within 11–20 points were considered ‘satisfactory’ (≥50%). Associations between the explanatory and outcome variables were subjected to likelihood stepwise backward multivariable logistic regression models to control for confounding and test for effect modification. A p<0.05 was considered statistically significant in all analyses.

## Results

### Distribution of camels and sero-prevalence for RVF Antibodies

A total of 720 camels were sampled from the two states; 420 were sampled from Jigawa and 300 from Katsina States, respectively (Table 1). An overall prevalence of 19.9 % (95 % CI 17.07-22.90) was obtained. From the sampled camels, a total of 83 (19.8%) camels were positive for RVFV antibodies in Jigawa State, while 60 (20%) were positive in Katsina State. Camels sampled from Katsina State showed a slightly higher prevalence rate compare to those in Jigawa State. Among the camels sampled from the seven Local Government Areas in Jigawa and Katsina States, camels from SuleTankarkar LGA had the highest seropositivity (33.3%) while the lowest seropositivity (3.30%) was recorded from camels in Daura LGA (Table 1).

**Table 1:** Demographic Distribution of sampled Camels and Prevalence of RVF virus Antibodies in Jigawa and Katsina States, Nigeria

| Category | Total no. sampled | No .positive on IgG | Percentage (%) | 95% Confidence interval |
| --- | --- | --- | --- | --- |

| **State** | **Location** |  |  |  |  |
| --- | --- | --- | --- | --- | --- |
|  | Jigawa | 420 | 83 | 19.8 | 16.16- 23.78 |
|  | Katsina | 300 | 60 | 20.0 | 15.76- 24.28 |
|  | **Total** | **720** | **143** | **19.9** | **17.07-22.90** |
| **Jigawa** | Maigatari | 220 | 37 | 16.8 | 12.31-22.20 |
|  | Babura | 100 | 18 | 18.0 | 11.38-26.45 |
|  | SuleTankarkar | 60 | 20 | 33.3 | 22.31-45.93 |
|  | Gumel | 40 | 8 | 20.0 | 9.75- 34.47 |
| **Katsina** | Maiadua | 150 | 37 | 24.7 | 18.27-32.04 |
|  | Jibiya | 90 | 21 | 23.3 | 15.47-32.89 |
|  | Daura | 60 | 2 | 3.3 | 0.56-10.58 |
|  | **Total** | **720** | **143** | **19.9** | **17.07-22.90** |
| **Sex** |  |  |  |  |  |
|  | Male | 475 | 93 | 19.6 | 16.20- 23.33 |
|  | Female | 245 | 50 | 20.4 | 15.71- 25.80 |
|  | **Total** | **720** | **143** | **19.9** | **17.07- 22.90** |
| **Age** |  |  |  |  |  |
|  | **Age group** |  |  |  |  |
|  | 0- 5 years | 160 | 32 | 20.0 | 14.34-26.73 |
|  | 6 - 10 years | 368 | 77 | 20.9 | 17.00-25.31 |
|  | 11 - 15 years | 189 | 34 | 18.0 | 13.00-23.95 |
|  | 16 - 20 years | 3 | 0 | 0.0 | - |
|  | **Total** | **720** | **143** | **19.9** | **17.07- 22.90** |

A total of 475 male and 245 female camels were sampled. Of the 475 males, 93 (19.6%) were positive, and females 50 (20.4%) of the 245 sampled were positive, with the female recording a slightly high seropositivity. All camels sampled were categorized into four (4) age groups. Camel between the age group 6-10 years recorded the highest prevalence (20.9%). Sex, age, and location were considered as risk factors and analyzed using multivariate regression with age and sex having no statistical significance however Sule Tankarkar ( OR 2.47; 95%CI 1.31-4.72; p=0.007) in Jigawa State and Maidua ( OR 1.62; 95%CI 0.97-2.73 p= 0.030) Katsina State were statistical significance on the occurrence of RVF in the study area (Table 2).

**Table 2:**
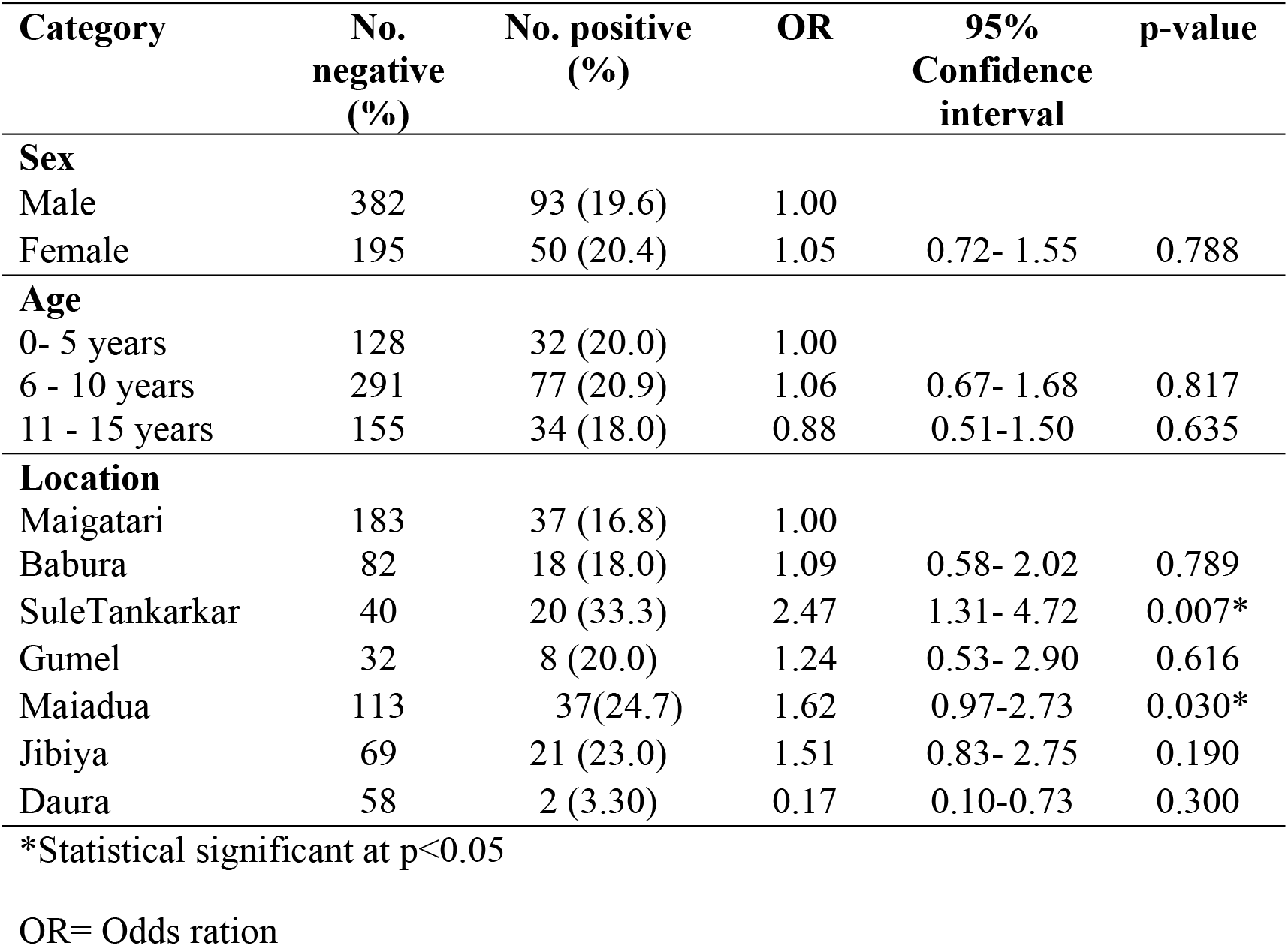
Multivariate analysis of risk factors associated with RVF sero-positivity in camels

| Category | No.<br>negative<br>(%) | No. positive<br>(%) | OR | 95%<br>Confidence<br>interval | p-value |
| --- | --- | --- | --- | --- | --- |
| **Sex** |  |  |  |  |  |
| Male | 382 | 93 (19.6) | 1.00 |  |  |
| Female | 195 | 50 (20.4) | 1.05 | 0.72- 1.55 | 0.788 |
| **Age** |  |  |  |  |  |
| 0- 5 years | 128 | 32 (20.0) | 1.00 |  |  |
| 6 - 10 years | 291 | 77 (20.9) | 1.06 | 0.67- 1.68 | 0.817 |
| 11 - 15 years | 155 | 34 (18.0) | 0.88 | 0.51-1.50 | 0.635 |
| **Location** |  |  |  |  |  |
| Maigatari | 183 | 37 (16.8) | 1.00 |  |  |
| Babura | 82 | 18 (18.0) | 1.09 | 0.58- 2.02 | 0.789 |
| SuleTankarkar | 40 | 20 (33.3) | 2.47 | 1.31- 4.72 | 0.007* |
| Gumel | 32 | 8 (20.0) | 1.24 | 0.53- 2.90 | 0.616 |
| Maiadua | 113 | 37(24.7) | 1.62 | 0.97-2.73 | 0.030* |
| Jibiya | 69 | 21 (23.0) | 1.51 | 0.83- 2.75 | 0.190 |
| Daura | 58 | 2 (3.30) | 0.17 | 0.10-0.73 | 0.300 |

\*Statistical significant at $p < 0.05$
OR= Odds ration

Results of the seropositivity were matched with coordinates of the sampled local government areas captured using the global positioning system (Fig 2) and imported ArcGIS Version 9.3. to produce a disease risk map (Fig 3).

**Fig 2:**
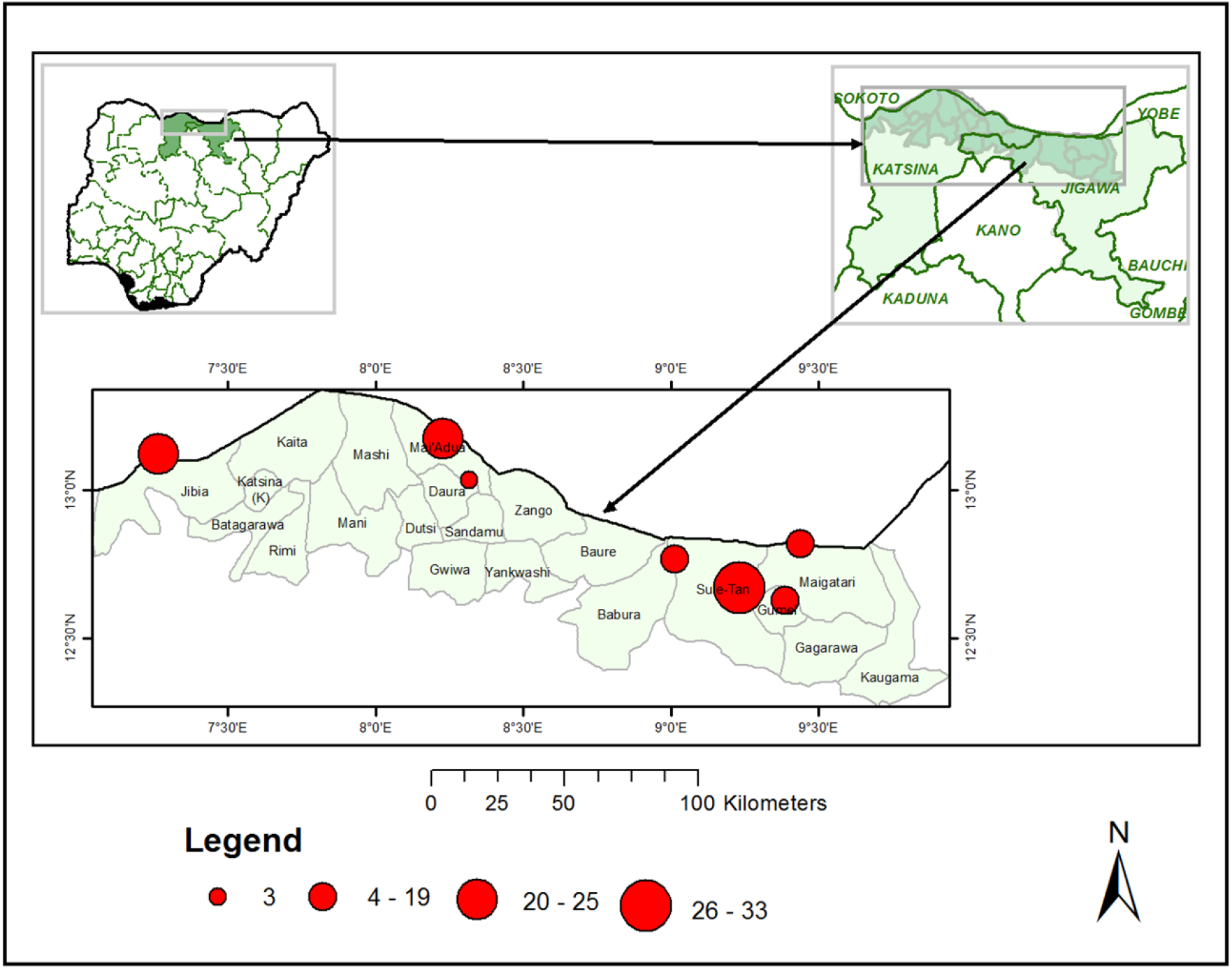
Map of Nigeria top left, Sampled States top right and study areas with seropositivity shown in red circles captured using global positioning system.

**Fig 3:**
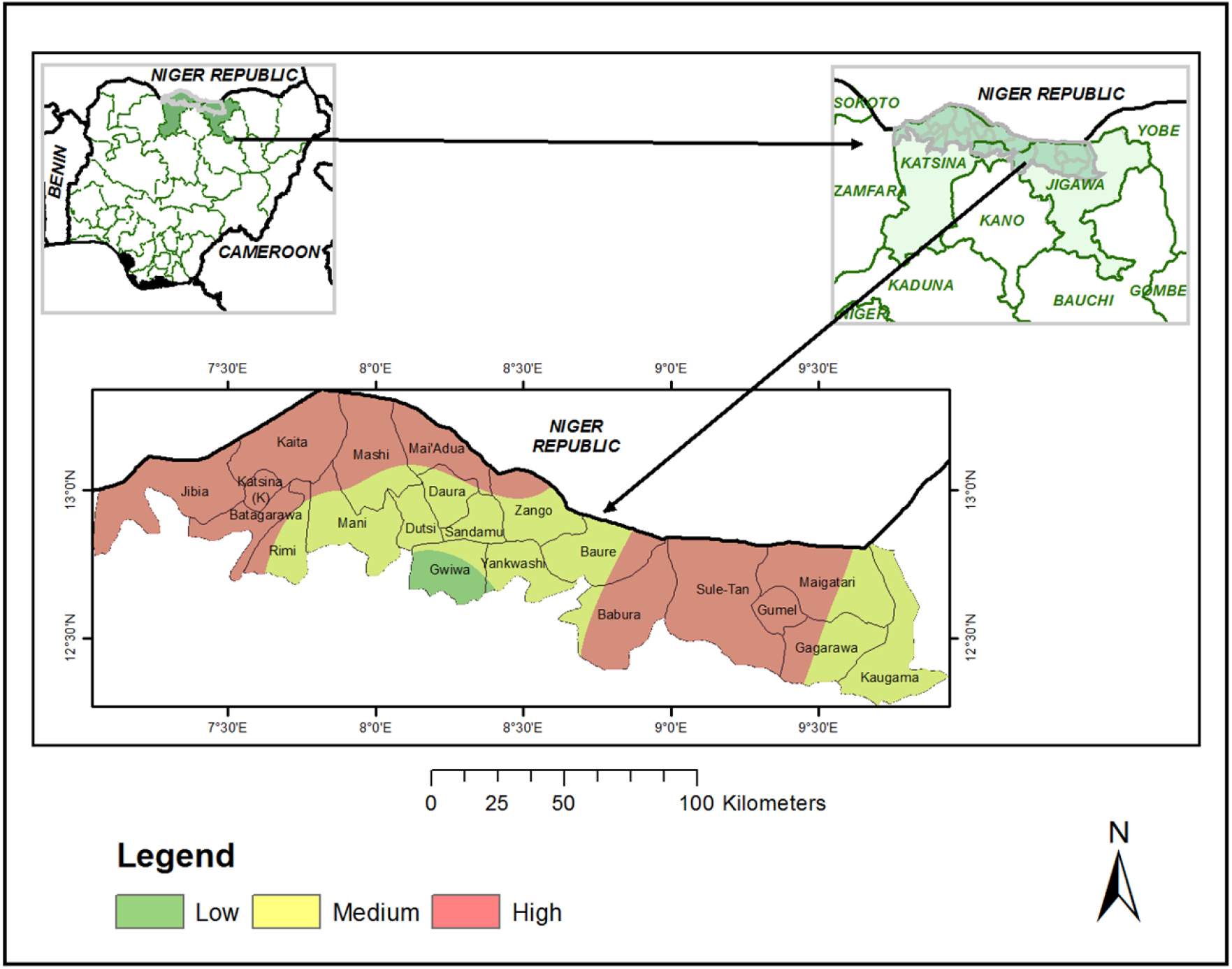
Risk Map of the study areas indicating the level of occurrence of RVFV. Green color indicates low risk, yellow indicates medium risk and red high risk.

### Socio-demographic characteristics of responding camel owners

Structured questionnaires were administered to 88 camels to respondents. Their ages were grouped into six from 20-29, 30-39, 40-49, 50-59, 60-69, and 70-79, with the highest respondents being from age group 40-49 (n=24/88) which is 27.3%. On the level of education, 65.9% (n=58/88) had no formal education. Also, their occupation was grouped into agro-pastoralists and transhumance, or nomadic pastoralists were more 56.8% (n=50/88). All the respondents were males, 100% (n=88) (Table 3).

**Table 3.**
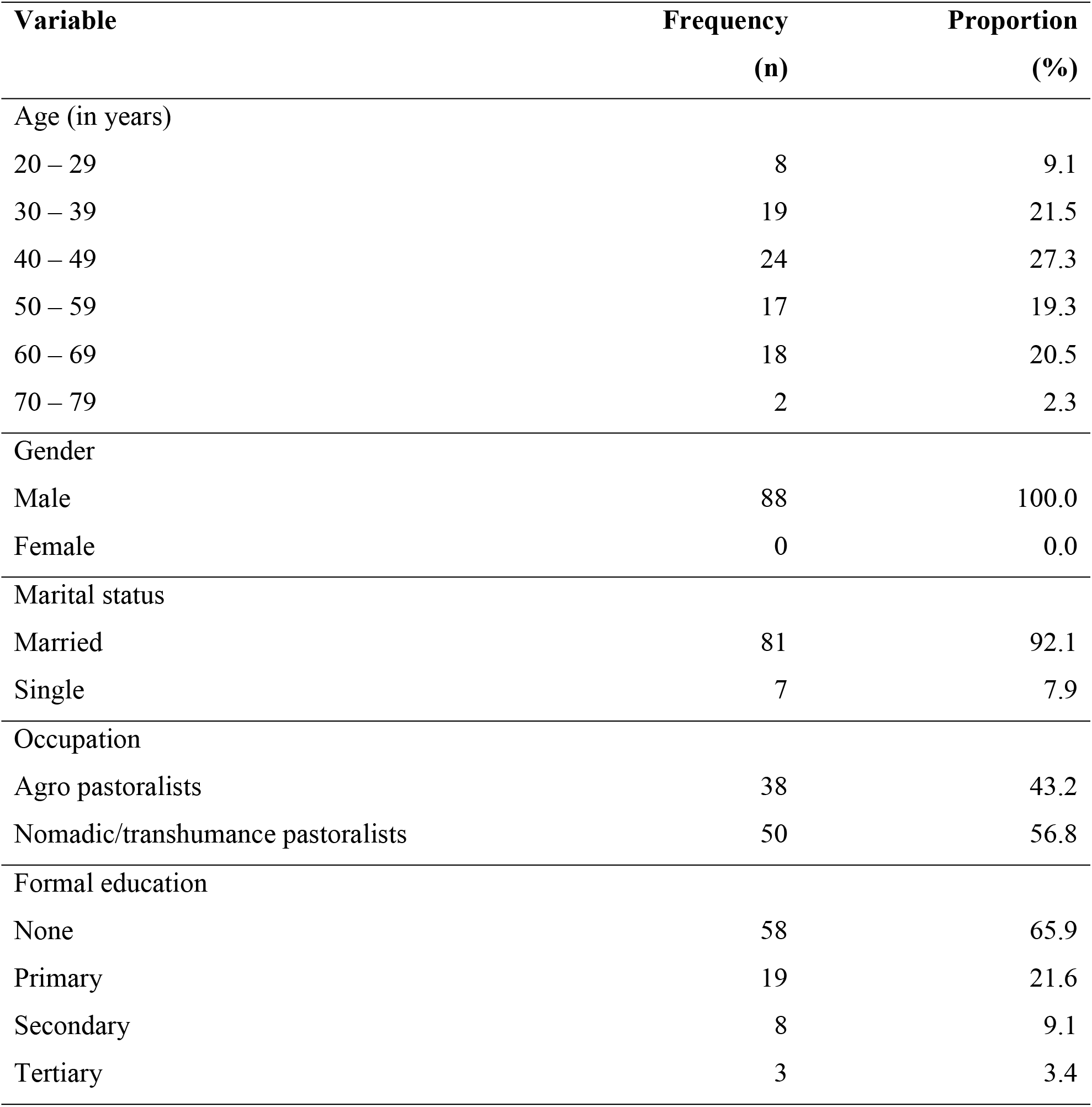
Socio-demographic characteristics of responding camel owners in North-West Nigeria.

### Camel owners’ knowledge about Rift Valley fever in camels in North-west Nigeria

Respondents were assessed on some clinical signs associated with RVF, and abortion in pregnant camels was the highest with 33% (n= 29/88) (95% CI 23.75-43.26). Only 19.3% (n=17/88) (95% CI 12.07-28.56) were known about the zoonotic implication of RVF. We sought to know about the endemicity of RVF in their environment, and 28.4% (n= 25/88) responded in the affirmative (Table 4).

**Table 4.** Camel owners’ knowledge about Rift Valley fever in North-west Nigeria

| Variable | Yes<br>(n) | Proportion<br>(%) | 95% Confidence<br>interval |
| --- | --- | --- | --- |
| Sign of RVF in camel |  |  |  |
| Anorexia | 22 | 25.0 | 16.8-34.82 |
| Listlessness in newborn calves | 12 | 13.6 | 7.61- 22.03 |
| Sudden deaths in newborns | 18 | 20.5 | 13.0- 29.83 |
| Abortions in pregnant animals | 29 | 33.0 | 23.75- 43.26 |
| Hemorrhages | 15 | 17.1 | 10.25- 25.98 |
| Fetid diarrhea | 26 | 29.6 | 20.73- 39.69 |
| All of the above | 5 | 5.7 | 2.11- 12.14 |
| Zoonotic nature of RVF |  |  |  |
| RVF can be transmitted from camels<br>to humans | 17 | 19.3 | 12.07- 28.56 |
| Endemic nature |  |  |  |
| RVF is endemic in the area | 25 | 28.4 | 19.74- 38.48 |

### Mitigation measures practice against Rift Valley fever in camel settlements in North-west Nigeria

On the mitigation measures practiced against RVF in camel settlements in the study areas, avoiding the culture of animal loaning, borrowing or dowry ranked top with 72.7% (n= 64/88) while use of repellents on animal against arthropods 17.1%(15/88) and avoiding ponds or swampy areas during grazing 17.1% (n=15/88) were the least (Table 5).

**Table 5.**
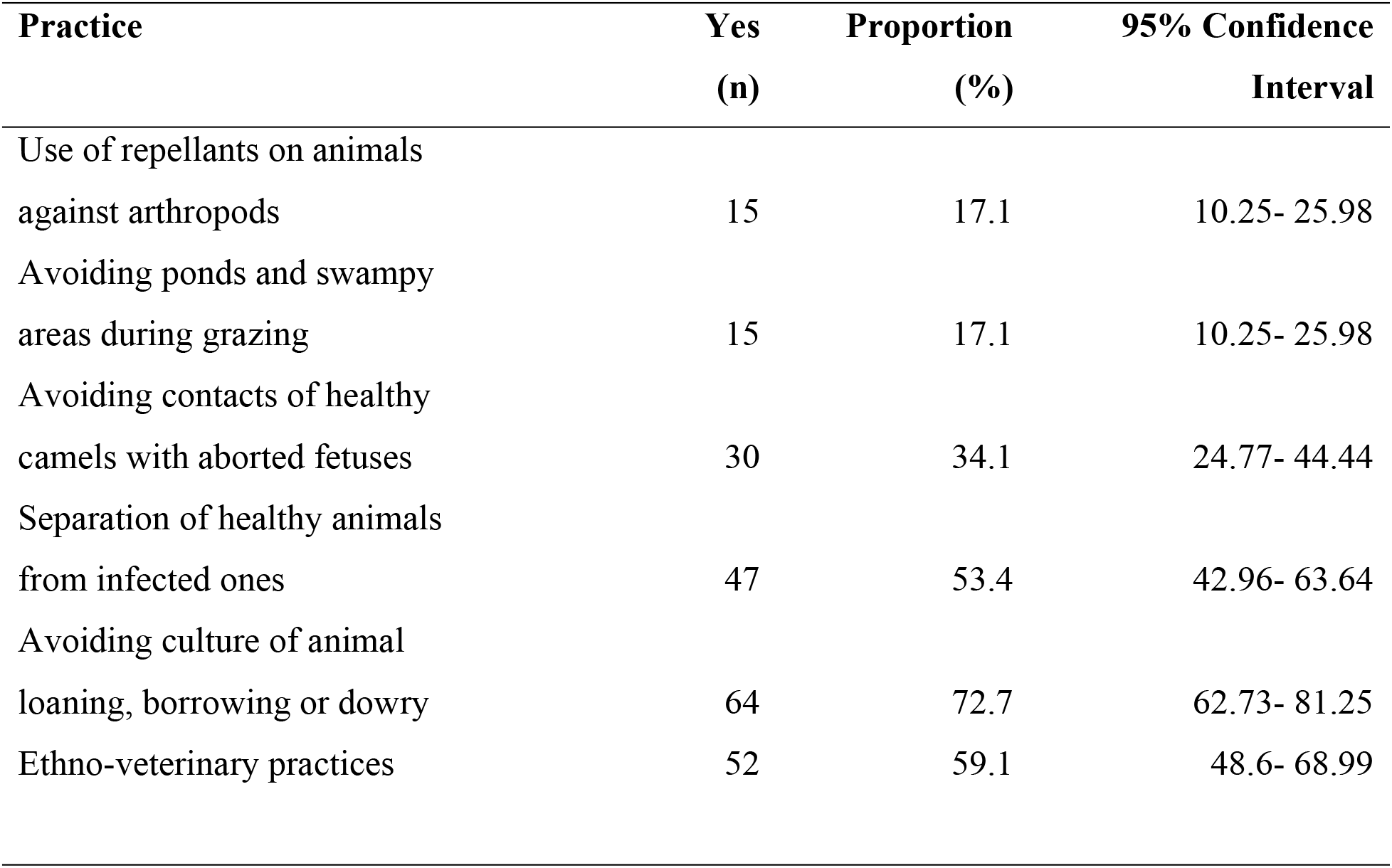
Mitigation measures practice against Rift Valley fever in camel settlements in North-west Nigeria

| Practice | Yes<br>(n) | Proportion<br>(%) | 95% Confidence<br>Interval |
| --- | --- | --- | --- |
| Use of repellants on animals<br>against arthropods | 15 | 17.1 | 10.25- 25.98 |
| Avoiding ponds and swampy<br>areas during grazing | 15 | 17.1 | 10.25- 25.98 |
| Avoiding contacts of healthy<br>camels with aborted fetuses | 30 | 34.1 | 24.77- 44.44 |
| Separation of healthy animals<br>from infected ones | 47 | 53.4 | 42.96- 63.64 |
| Avoiding culture of animal<br>loaning, borrowing or dowry | 64 | 72.7 | 62.73- 81.25 |
| Ethno-veterinary practices | 52 | 59.1 | 48.6- 68.99 |

### Multivariable logistic regressions of environmental factors that influence Rift Valley fever occurrence in camel settlements in North-west Nigeria

Environmental risk factors for the occurrence of RVF were assessed using multivariable logistic regression between agro-pastoralists and nomadic pastoralists (Table 6). High rainfall was the only factor that showed no significance among nomadic pastoralists settlements despite odds OR 2.2 (95%CI 0.93-5.20; p=0.070).

**Table 6.**
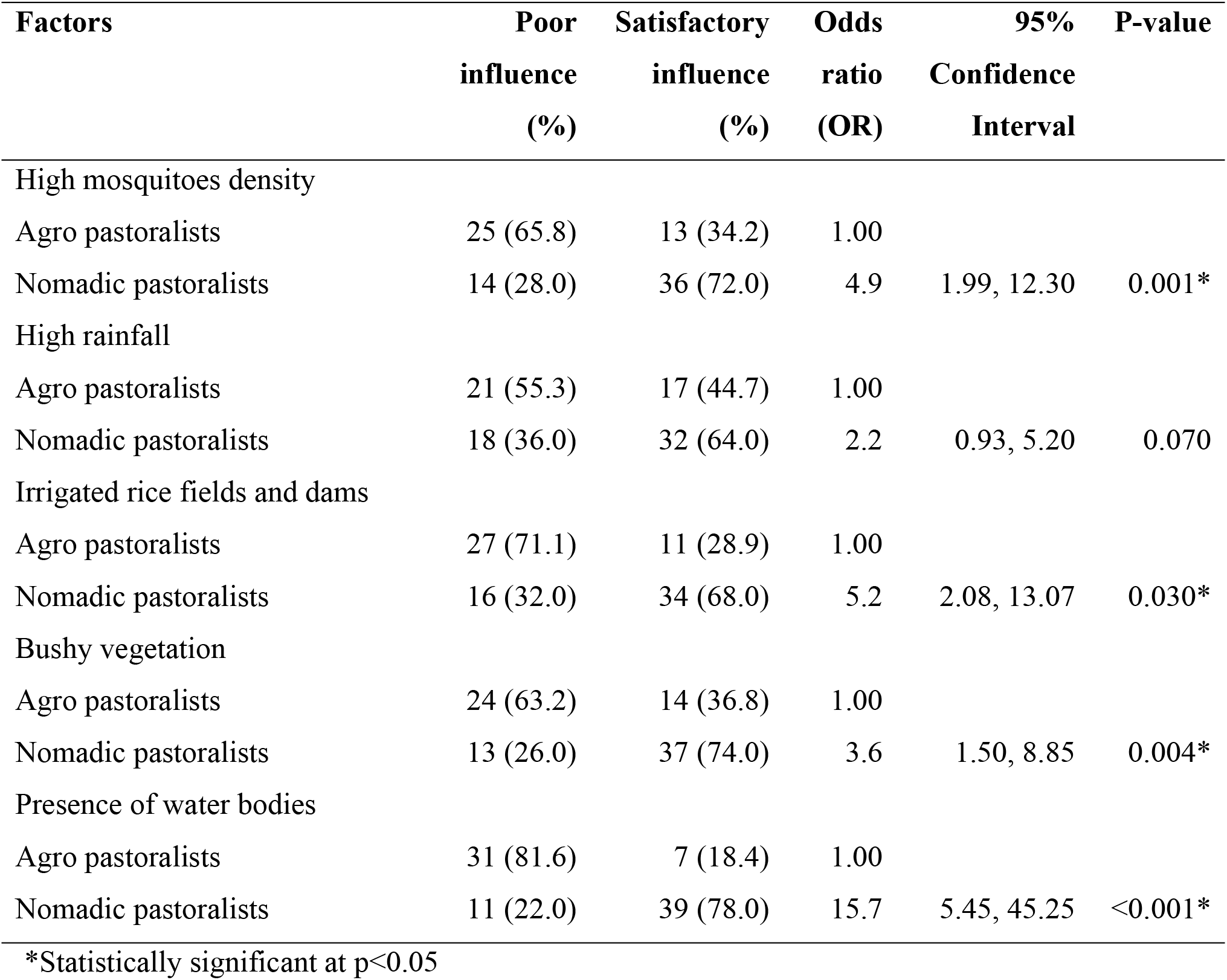
Multivariable logistic regressions models of environmental precipitating factors that influence Rift Valley fever occurrence in camel settlements in North-west Nigeria

## Discussion

The result of this study demonstrated the prevalence of RVFV- specific immunoglobulin G (IgG) antibodies using a competitive enzyme-linked immunosorbent assay (c-ELISA) in one-humped camels (*Camelus dromedarius*) in Jigawa and Katsina States of Nigeria despite the absence of outbreak which indicates the cryptic circulation of RVFV in the camel population. The study revealed an overall prevalence of 19.9% among sampled camels from the two States. Katsina State recorded a higher prevalence of 20% while Jigawa State had 19.8%. However, these differences were not statistically significant. In Nigeria, [18], recorded a prevalence of 3.03% in camels.[19] reported a slight increase of prevalence of 3.13% when compared to the present study, there appears to be a remarkable increase in the prevalence of Rift Valley fever which could be due to increased movement of camels mainly due to transhumance from Nigeria to neighbouring countries and back to Nigeria. There is also the possibility of increased vector populations and seasonal changes due to global warming and climate change with increased vectors, which could serve as mechanical and biological vectors in the amplification of RVFV and may maintain the virus at low level. Increased mobility of camel owners due to insecurity in the Sahel as the camel owners attempt to migrate to safe areas, they may get exposed to several precipitating factors of RVF.

In comparison to other studies the prevalence of 19.9% is however lower than previous surveys of Rift Valley fever in camels reported in Niger Republic 47.5% [20], 45% in Mauritania [21], and 38.5% in Tanzania [22]. The higher prevalence reported could be attributed to the epidemics of RVF reported in these affected regions. In Sudan, [23] reported a prevalence of 9.6% in their studies and for the first time in Turkey, RVFV antibodies were reported in camels, with a prevalence of 1.3% [24]. Again, the areas in Sudan and Turkey had never experienced any epizootic of RVFV in livestock; however, antibodies were demonstrated, which may be an indication for a covert circulation waiting for the right environment for an outbreak to take place.

The prevalence of RVFV antibodies recorded in Katsina State may be attributed to the fact that it shares an international border with Zinder and Maradi Provinces of Niger Republic. Camels from Nigeria migrate into the neighboring country (Niger) freely at the onset of the rainy season (May-June) and return at the start of dry season (October-November) without being monitored at the borders. Maradi Province in Niger Republic which shares border with Katsina State is located South East of Tahoua Province in Niger Republic which reported an outbreak of RVF between August and September 2016 with over 300 fatalities in cattle, goats, sheep and camels and more than 33 deaths in humans [25]. Most of the affected people were pastoralists who had participated in an annual festival at the border between Niger Republic and Burkina Faso and Mali and had direct contact with infected animal tissues or body fluids [26].

Also, the high prevalence in camels recorded as compared to other studies in camels in Nigeria could be due to increased construction of irrigation dams and agricultural schemes, which may have contributed to suitable climatic condition for the survival and proliferation of *Aedes* and *Culex* which are significant vectors in the transmission of RVFV. [27] reported that vector-borne diseases are always associated with the geographical areas where these vectors are found, and given the right environmental conditions, arboviruses can persist. The increased positivity could be because there has been increased demand for camel meat and products (such as the milk and urine) due to increased Nigerian population and also the relatively cheap cost as compare to beef. [28] observed that since the outbreak of Rinderpest in Nigeria in the 1980s, camel meat consumption has increased dramatically.

Prevalence based on location, the highest seropositivity recorded in Sule Tankarkar could be attributed to the presence of two transhumance routes that transverse the local government areas. Also, The proximity of the local government area to (Zinder Province) Niger Republic where it covers a relatively large area coupled with unrestricted movement of stray camels into Jigawa State as reported by the Jigawa state government in 2017 and more recently in 2019 by Federal Agencies (Federal Road Safety Corps and Nigerian Immigration Service) that over 7000 strayed camels from Sudan, Mali, Burkina Faso, Tchad and Niger republic were found in Jigawa State [29,30] may have contributed to the high prevalence recorded. This corroborates with the studies of[31], who observed that the introduction of RVFV into Egypt was attributed to porous borders where camels come from Sudan and Ethiopia enter Egypt either legally or otherwise. Since an outbreak of RVFV had occurred in 2016 in the neighbouring Niger Republic and coupled with the presence of an international livestock market in a neighbouring Maigatari local government area in Jigawa State where different animals from differentAfrican countries and various states conglomerate for sale, it may be possible that movement of some infected camels from these countries contributed to the high seropositivity recorded. [27,32] had all observed that given a favourable condition, transboundary diseases have the potential to spread over a considerable distance.

Similarly, Mai’adua local government area in Katsina State recorded the highest prevalence of 24.7%. It was observed that camels from Mai’adua local government area are approximately two times more likely to be infected with RVF (OR= 1.67). This could be attributed to the boundary shared with Niger republic; presence of an international market which may increase contact with animals that have come from the neighbouring countries with the infection and the presence of hematophagous insects like *Hippobosca equiinus, Hyalomma dromederii, Stomoxys, Amblyomma variegatum, Argas persicus and ornithodorous* in this area which feed on these animals and become new vectors that amplify the spread of RVFV [33].

[31], [34] and [35] observed that importation of ruminants was a significant source of Rift Valley fever virus and other pathogens introduced into places like Egypt, Saudi Arabia, and Yemen which recorded outbreaks in 2000. Lastly, the presence of Sabke Dam located in Maiadua LGA could be one of the precipitating factor for the high prevalence observed in this study due to accessibility of this Dam by various livestock species for source of water and also increases contact with animals and hematophagous insects that may be found around the area. Meanwhile,[36] and [37] reported an epizootic of RVF after the construction Aswan Dam in Egypt in 1977. Similarly, the presence of the Diama Dam in Mauritania contributed to the first outbreak encountered in Mauritania in 1987 [38–39].

Daura is a semi-urban settlement in which most camel owners practice semi-intensive systems of rearing as most are owned by the royal family who use them during festivals and for prestige. This could suggest the low prevalence recorded in this area. [40] observed that urban settlement is unfavourable to the vector species, but also due to limited livestock movements, there is little interaction between neighbouring herds, thus the possibility of limited infection.

Higher seropositivity, though with no significant difference, was recorded in females than males. It is a general observation that female animals stay longer than males in the herd. Therefore the longer the females stay in the herd, the longer they are exposed to risk factors associated with RVF [23]. This corroborates with the report by [41] in Mauritania and [40] in Tanzania regarding significant sero-positivity with female sex.

Based on age, camels within the 6-10 age category had the highest prevalence. This age group is considered the age of maturity or adulthood for camels when they are released into the pasture for grazing and travel long distances where they may be exposed to infected mosquitoes and other hematophagous insects and hence the high positivity among this age group. This agrees with some authors, who all reported higher risks of older camels to Rift Valley fever infection like [22–23,32]. The camel owners give younger camels more attention and provide them with the necessary care which minimizes the risk of infectious diseases contraction like vector-borne diseases due to short distances covered.

Transhumance (nomadic) pastoralists were likely to be affected by environmental risk factors on the occurrence of Rift Valley fever in both Jigawa and Katsina States. In this study, an assessment was carried out among camel owners on some environmental risk factors such as high mosquito density, irrigated rice fields and dams, bushy vegetation and presence of water bodies in both Katsina and Jigawa States and were significant for the occurrence of Rift Valley fever in the study areas with the exception of high rainfall. The significance of these assessments could be attributed to the movements of pastoralists strategically to different countries in search of ecologically viable resources for their animals and therefore, being exposed to these risk factors. These observations agree with the works of [42], where they observed that RVF was more likely to occur in densely bushed areas. Similarly, [43] observed that in arid and semi-arid areas where irrigations are practiced to mitigate the scourge of food insecurity challenges, the establishment of dams and large pools of water maintains high humidity thereby increasing the suitability of RVF endemicity. On the non-significance of high rainfall to the occurrence of RVF, this study was carried out in arid and semi-arid areas of Jigawa and Katsina States with low yearly rainfall, which makes it a conducive for camel production. This corroborates with the works of[44], who observed that camel is a desert ship that does not tolerate the environment with high rainfall for its survival. Also, some authors have identified low rainfall as one of the ecological factors necessary for the occurrence of RVF in an inter-epizootic periods [45,46].

In conclusion, though this study showed increased serological evidence of the presence of RVFV in camels in Nigeria, there is need for nationwide surveillance on serological and virological status of the RVFV in livestock, vectors and human population as this will provide a base line data for better tracking and forecasting of RVF outbreak. Also, bordered local government areas with Niger Republic are at high risk of contracting RVF, there is a need for Nigerian governments to have quarantine units across borders for screening animals coming from neighboring countries for transboundary infectious diseases such as RVF.

## Acknowledgments

Authors wish to thank the Directors of Veterinary Services in Jigawa and Katsina States for their cooperation in carrying out this research in their respective states. Special thanks to Habila Iliyasu and Salisu Usman, Sanusi Faby, Zaradeen Adamu, and Malam Sani Abdulmumuni for their assistance during the sample collection.

## References

1. Davies FG, Martin V. Recognizing rift valley fever. Vet Ital. 2006 Jan;42(1):31–53.

2. . Adams MJ, Lefkowitz EJ, King AM, Harrach B, Harrison RL, Knowles NJ, Kropinski AM, Krupovic M, Kuhn JH, Mushegian AR, Nibert M. Changes to taxonomy and the International Code of Virus Classification and Nomenclature ratified by the International Committee on Taxonomy of Viruses (2017). Archives of virology. 2017 Aug 1;162(8):2505–38.

3. Schmaljohn CS. Bunyaviridae: the viruses and their replication. Fields of Virology. 2001.

4. Rich KM, Wanyoike F. An assessment of the regional and national socio-economic impacts of the 2007 Rift Valley fever outbreak in Kenya. The American journal of tropical medicine and hygiene. 2010 Aug 5;83(2_Suppl):52–7.

5. Chengula AA, Mdegela RH, Kasanga CJ. Socio-economic impact of Rift Valley fever to pastoralists and agro pastoralists in Arusha, Manyara and Morogoro regions in Tanzania. Springerplus. 2013 Dec 1;2(1):549.

6. Mansfield KL, Banyard AC, McElhinney L, Johnson N, Horton DL, Hernández-Triana LM, Fooks AR. Rift Valley fever virus: A review of diagnosis and vaccination, and implications for emergence in Europe. Vaccine. 2015 Oct 13;33(42):5520–31.

7. Balkhy HH, Memish ZA. Rift Valley fever: an uninvited zoonosis in the Arabian peninsula. International journal of antimicrobial agents. 2003 Feb 1;21(2):153–7.

8. Jansen van Vuren P, Kgaladi J, Patharoo V, Ohaebosim P, Msimang V, Nyokong B, Paweska JT. Human Cases of Rift Valley Fever in South Africa, 2018. Vector-Borne and Zoonotic Diseases. 2018 Nov 28;18(12):713–5.

9. Chevalier V, Pépin M, Plee L, Lancelot R. Rift Valley fever-a threat for Europe?. Euro Surveillance. 2010;15(10):18–28.

10. Ferguson W. Identification of Rift Valley fever in Nigeria. Bull Epizoot Dis Afr.1959;7:317–8.

11. Lee VH. Isolation of viruses from field populations of Culicoides (Diptera: Ceratopogonidae) in Nigeria. Journal of Medical Entomology. 1979 Sep 12;16(1):76–9.

12. Pepin M, Bouloy M, Bird BH, Kemp A, Paweska J. Rift Valley fever virus (*Bunyaviridae: Phlebovirus*): an update on pathogenesis, molecular epidemiology, vectors, diagnostics and prevention. Veterinary research. 2010 Nov 1;41(6):61.

13. Meegan JM. The Rift Valley fever epizootic in Egypt 1977–1978 1. Description of the epizootic and virological studies. Transactions of the Royal Society of Tropical Medicine and Hygiene. 1979 Jan 1;73(6):618–23.

14. Baba M, Masiga DK, Sang R, Villinger J. Has Rift Valley fever virus evolved with increasing severity in human populations in East Africa? Emerging microbes & infections. 2016 Jan 1;5(1):1–0.

15. Lumley S, Horton D, Hernandez-Triana LL, Johnson N, Fooks AR, Hewson R. Rift Valley fever virus: strategies for maintenance, survival and vertical transmission in mosquitoes. Journal of General Virology. 2017 May 30;98(5):875–87.

16. Kortekaas J, Kant J, Vloet R, Cêtre-Sossah C, Marianneau P, Lacote S, Banyard AC, Jeffries C, Eiden M, Groschup M, Jäckel S. European ring trial to evaluate ELISAs for the diagnosis of infection with Rift Valley fever virus. Journal of virological methods. 2013 Jan 1;187(1):177–81.

17. Oloyede-Kosoko SO, Akingbogun AA. Geospatial Information in Public Health: Using Geographical Information System to Model the Spread of Tuberculosis. FIG Working Week, Environment for Sustainability. Richardson. 2013.

18. Ezeifeka GO, Umoh JU, Belino ED, Ezeokoli CD. A serological survey for Rift Valley fever antibody in food animals in Kaduna and Sokoto States of Nigeria. International journal of zoonoses. 1982 Dec;9(2):147–51.

19. Olaleye OD, Tomori O, Schmitz H. Rift Valley fever in Nigeria: infections in domestic animals. Revue scientifique et technique-Office international des épizooties. 1996 Sep 1;15:937–46.

20. Mariner JC, Morrill J, Ksiazek TG. Antibodies to hemorrhagic fever viruses in domestic livestock in Niger: Rift Valley fever and Crimean-Congo hemorrhagic fever. The American journal of tropical medicine and hygiene. 1995 Sep 1;53(3):217–21.

21. Jäckel S, Eiden M, El Mamy BO, Isselmou K, Vina‐Rodriguez A, Doumbia B, Groschup MH. Molecular and Serological Studies on the R ift V alley Fever Outbreak in M auritania in 2010. Transboundary and emerging diseases. 2013 Nov;60:31–9.

22. Swai ES, Sindato C. Seroprevalence of Rift Valley fever virus infection in camels (dromedaries) in northern Tanzania. Tropical animal health and production. 2015 Feb 1;47(2):347–52.

23. Abdallah MM, Adam IA, Abdalla TM, Abdelaziz SA, Ahmed ME, Aradaib IE. A survey of rift valley fever and associated risk factors among the one-humped camel (Camelus dromedaries) in Sudan. Irish veterinary journal. 2015 Dec;69(1):6.

24. Gür S, Kale M, Erol N, Yapici O, Mamak N, Yavru S. The first serological evidence for Rift Valley fever infection in the camel, goitered gazelle and Anatolian water buffaloes in Turkey. Tropical animal health and production. 2017 Oct 1;49(7):1531–5.

25. World Health Organization (2016). Rift Valley fever in Niger.http://www.who.int/csr/don/29-september-2016-rift-valley-fever-niger/en/. Retrieved on 8 Novemeber 2017

26. International Federation of Red Cross and Red Crescent Societies (2016). Niger Red Cross responds to Rift Valley fever outbreak. https://reliefweb.int/report/niger/niger-red-cross-responds-rift-valley-fever-outbreak. Retrieved March 27th 2017.

27. Martin V, Chevalier V, Ceccato P, Anyamba A, De Simone L, Lubroth J, de La Rocque S, Domenech J. The impact of climate change on the epidemiology and control of Rift Valley fever. Revue Scientifique et Technique-Office international des épizooties. 2008;27(2):413–26.

28. Abubakar UB, Mohammed FU, Shehu SA, Mustapha RA. Foetal wastage in camels slaughtered (Camelus dromedarius) at Maiduguri Abattoir, Borno State, Nigeria. International Journal of Tropical Medicine. 2010;5(4):86–8.

29. Hamagam, A.M (2017). Dutse.Over 7,000 stray camels enter Jigawa from Sudan, Mali. https://www.dailytrust.com.ng/news/general/over-7-000-stray-camels-enter-jigawa-from-sudan-mali/190639.html

30. Muhammad, K. (2019). Sixteen dead as foreigners ‘camels cause accidents on Jigawa roads. Retrieved from https://dailypost.ng/2019/04/22/16-dead-foreigners-camels-cause-accidents-jigawa-roads/

31. Napp S, Chevalier V, Busquets N, Calistri P, Casal J, Attia M, Elbassal R, Hosni H, Farrag H, Hassan N, Tawfik R. Understanding the legal trade of cattle and camels and the derived risk of Rift Valley Fever introduction into and transmission within Egypt. PLoS Neglected Ttropical Diseases. 2018 Jan 19;12(1):e0006143.

32. Di Nardo A, Rossi D, Saleh SM, Lejlifa SM, Hamdi SJ, Di Gennaro A, Savini G, Thrusfield MV. Evidence of Rift Valley fever seroprevalence in the Sahrawi semi-nomadic pastoralist system, Western Sahara. BMC Veterinary Research. 2014 Dec;10(1):92.

33. Linthicum KJ, Britch SC, Anyamba A. Rift Valley fever: an emerging mosquito-borne disease. Annual review of Entomology. 2016 Mar 11;61:395–415.

34. Kamal SA. Observations on rift valley fever virus and vaccines in Egypt. Virology journal. 2011 Dec;8(1):532.

35. Chu DK, Poon LL, Gomaa MM, Shehata MM, Perera RA, Zeid DA, El Rifay AS, Siu LY, Guan Y, Webby RJ, Ali MA. MERS coronaviruses in dromedary camels, Egypt. Emerging Infectious Diseases. 2014 Jun;20(6):1049.

36. Meegan JM. The Rift Valley fever epizootic in Egypt 1977–1978 1. Description of the epizootic and virological studies. Transactions of the Royal Society of Tropical Medicine and Hygiene. 1979 Jan 1;73(6):618–23.

37. Meegan JM, Hoogstraal H, Moussa MI. An epizootic of Rift Valley fever in Egypt in 1977. The Veterinary Record. 1979 Aug;105(6):124–5.

38. Digoutte JP, Peters CJ. General aspects of the 1987 Rift Valley fever epidemic in Mauritania. Research in Virology. 1989 Jan 1;140:27–30.

39. Jouan A, Coulibaly I, Adam F, Philippe B, Riou O, Leguenno B, Christie R, Merzoug NO, Ksiazek T, Digoutte JP. Analytical study of a Rift Valley fever epidemic. Research in Virology. 1989 Jan 1;140:175–86.

40. Sumaye RD, Geubbels E, Mbeyela E, Berkvens D. Inter-epidemic transmission of Rift Valley fever in livestock in the Kilombero River Valley, Tanzania: a cross-sectional survey. PLoS Neglected Tropical Diseases. 2013 Aug 8;7(8):e2356.

41. El Mamy AB, Baba MO, Barry Y, Isselmou K, Dia ML, Hampate B, Diallo MY, El Kory MO, Diop M, Lo MM, Thiongane Y. Unexpected rift valley fever outbreak, Northern Mauritania. Emerging Infectious Diseases. 2011 Oct;17(10):1894.

42. Hightower A, Kinkade C, Nguku PM, Anyangu A, Mutonga D, Omolo J, Njenga MK, Feikin DR, Schnabel D, Ombok M, Breiman RF. Relationship of climate, geography, and geology to the incidence of Rift Valley fever in Kenya during the 2006–2007 outbreak. The American Journal of Tropical Medicine and Hygiene. 2012 Feb 1;86(2):373–80.

43. Mbotha D, Bett B, Kairu‐Wanyoike S, Grace D, Kihara A, Wainaina M, Hoppenheit A, Clausen PH, Lindahl J. Inter‐epidemic Rift Valley fever virus seroconversions in an irrigation scheme in Bura, south‐east Kenya. Transboundary and Emerging diseases. 2018 Feb;65(1):e55–62.

44. Jaji AZ, Elelu N, Mahre MB, Jaji K, Mohammed LG, Likita MA, Kigir ES, Onwuama KT, Saidu AS. Herd growth parameters and constraints of camel rearing in Northeastern Nigeria. Pastoralism. 2017 Dec;7(1):16.

45. Munyua PM, Murithi RM, Ithondeka P, Hightower A, Thumbi SM, Anyangu SA, Kiplimo J, Bett B, Vrieling A, Breiman RF, Njenga MK. Predictive factors and risk mapping for Rift Valley fever epidemics in Kenya. PLoS One. 2016 Jan 25;11(1):e0144570.

46. Njenga MK, Bett B. Rift Valley fever virus—how and where virus is maintained during inter-epidemic periods. Current Clinical Microbiology Reports. 2019 Mar 15;6(1):18–24.

